# Expanding the RNA polymerase biocatalyst solution space for mRNA manufacture

**DOI:** 10.1101/2024.01.28.577649

**Authors:** Edward Curry, Svetlana Sedelnikova, John Rafferty, Martyn Hulley, Adam Brown

## Abstract

All mRNA products are currently manufactured in *in vitro* transcription (IVT) reactions that utilize single-subunit RNA polymerase (RNAP) biocatalysts. Although it is known that discrete polymerases exhibit highly variable bioproduction phenotypes, including different relative processivity rates and impurity generation profiles, only a handful of enzymes are generally available for mRNA biosynthesis. This limited RNAP toolbox restricts strategies to design and troubleshoot new mRNA manufacturing processes, which is particularly undesirable given the continuing diversification of mRNA product lines towards larger and more complex molecules. Herein, we describe development of a high-throughput RNAP screening platform, comprising complementary *in silico* and *in vitro* testing modules, that enables functional characterisation of large enzyme libraries. Utilizing this system, we identified eight novel sequence-diverse RNAPs, with associated active cognate promoters, and subsequently validated their performance as recombinant enzymes in IVT-based mRNA production processes. By increasing the number of available characterized functional RNAPs by > 130% and providing a platform to rapidly identify further potentially useful enzymes, this work significantly expands the RNAP biocatalyst solution space for mRNA manufacture, thereby enhancing capability to achieve application and molecule-specific optimisation of product yield and quality.

## Introduction

The clinical success of SARS-Cov-2 vaccines established synthetic mRNA as an effective drug format, paving the way for hundreds of new mRNA-based vaccines and gene therapies to enter clinical trials (Qin et al., 2022; Webb et al., 2022). This has resulted in a sharp increase in global demand for mRNA production, and shifted mRNA manufacturing from a relatively niche process to one that underpins current and future strategies to treat monogenic disorders, cancer, and infectious diseases (Al Fayez et al., 2023; Liu et al., 2023; Vavilis et al., 2023). All such mRNA products are currently produced in standardised *in vitro* transcription (IVT) systems using single-subunit DNA-dependent phage RNA polymerase (RNAP) biocatalysts. While Salmonella phage SP6 and Enterobacteria phage T3 RNAPs can be utilised in certain contexts, the dominant biocatalyst choice for synthetic mRNA manufacture is the *Enterobacteria* phage T7 RNAP. This enzyme has undergone extensive protein engineering to improve bioproduction performance, predominantly via strategies to reduce formation of immunogenic product-related impurities such as short-abortive transcripts (Guilleres et al., 2005; Lyon and Gopalan, 2018) and double-stranded RNA species (Dousis et al., 2023; Wu et al., 2020).

Although T7 RNAP typically generates high product yields and acceptable product quality profiles, it is highly unlikely that a single one-size-fits-all biocatalyst approach will be optimal for all mRNA manufacturing contexts. Indeed, other bioproduction processes rely on biocatalyst toolbox approaches, such as the wide range of evolved and engineered Chinese Hamster Ovary cell factories utilised for recombinant protein manufacture (Fischer et al., 2015). The current unavailability of such an RNAP toolbox for mRNA IVT platforms restricts i) bioprocess design strategies, such as optimising temperature set-point to achieve quality target product profiles, and ii) molecule/application specific optimisation of product yield and quality. The latter is particularly pertinent given that mRNA product lines are diversifying to include larger and more complex molecular formats that present new biomanufacturing challenges, such as circular RNA (Bai et al., 2023; Qu et al., 2022; Zhu et al., 2022), long self-amplifying RNA transcripts (Blakney et al., 2021; Pourseif, 2022), and linear mRNA incorporating novel cap structures and modified nucleotides (Chen et al., 2022). Indeed, it should be anticipated that mRNA will follow the path of protein therapeutics, where product designers rapidly progressed from relatively simple molecules such as Insulin to highly-engineered formats (e.g. tri-specific antibodies) that require product-specific biocatalyst solutions (Tihanyi and Nyitray, 2020).

The available RNAP toolbox has recently been expanded by studies focussed on identifying and characterising individual enzymes with putative desirable bioproduction phenotypes. KP34 enhances 3’ homogeneity of product molecules (Lu et al., 2019), VSW-3 reduces doubled stranded RNA (dsRNA) impurities (Xia et al., 2022), and Syn5 exhibits increased processivity (Zhu et al., 2013), as compared to that achieved with T7. While these hypothesis-driven approaches have successfully identified new biocatalysts with novel functionalities, only six characterized RNAPs are currently publicly available for mRNA manufacture (although we note that some additional unpublished enzymes may be utilised in industrial settings). Accordingly, mRNA manufacturing solution spaces are severely limited, and, moreover, currently utilised ‘standardised’ biocatalysts such as T7 may have relatively poor performance characteristics (E.g. processivity, impurity generation) relative to the hundreds of ‘untested’ single-subunit phage RNAPs found in nature.

In this study, we address the paucity of biocatalysts available for IVT-based mRNA production. Using a combination of *in silico* and *in vitro* analyses we identify and functionally validate eight new sequence-diverse RNAPs, more than doubling the number of previously described enzymes for mRNA manufacture. In doing so, we describe development of a high-throughput screening system that can be utilised to rapidly select and test future RNAP libraries, facilitating further expansion of the biocatalyst solution space. Provision of a substantially expanded RNAP toolkit enhances capabilities to design and troubleshoot new molecule-specific manufacturing processes, which will be particularly useful for optimising yield and quality of complex next-generation mRNA products.

## 2. Materials and Methods

### 2.1 RNAP library creation

A starting library of 351 predicted RNAP sequences was collated from Uniprot, comprising all sequences annotated as predicted phage DNA-directed RNA polymerases. RNAPs were clustered by grouping RNAPs sharing sequence identity >85%, using the Clustal Omega online alignment tool (Madeira et al., 2019). RNAPs were further clustered using the MMSEQ2 online server, with a minimum sequence identity threshold of 85%, and coverage threshold of 70% (Steinegger and Söding, 2017). A representative RNAP from each cluster was chosen by totalling the matrix identity score to determine which polymerase in each cluster was most divergent in sequence to all others. Promoters for remaining RNAPs were predicted by PHIRE (Phage *in silico* regulatory elements) (Lavigne et al., 2004) using parameters of string lengths – 20, window size – 30, and degeneracy – 4. Predicted promoter sequences were verified with the *PhagePromoter* tool (Sampaio et al., 2019), with a probability threshold of 0.5.

### 2.2 Plasmid Construction

For coupled transcription-translation assay plasmids, RNAP sequences were synthesised and cloned into XhoI and XbaI restriction sites on the pTNT vector (Promega). To create the corresponding transcription templates, the NanoLuc gene (Promega) was cloned into XhoI and XbaI restriction sites in pTNT, before site directed mutagenesis to substitute the SP6 and T7 promoter with the promoter of interest. For RNAP overexpression plasmids, RNAP sequences were inserted between NdeI and XhoI sites on pET-29b (Novagen). Transcription templates were made by site directed mutagenesis to substitute the T7 promoter with the promoter of interest on pCMV-Cluc2 (New England Biolabs).

### 2.3 Cell free coupled transcription-translation assays

Coupled transcription-translation assays were carried out using the TNT SP6 Quick Coupled transcription/translation system (Promega). Reactions were assembled containing 8 μl TnT Quick Master Mix, 1 μl RNAP plasmid (40 ng/μl), 1 μl NanoLuc plasmid (80 ng/μl), and 0.2 μl 1 mM methionine. The assay proceeded at 30C for 1 hour. Samples were then diluted 500 fold in nuclease free water, and added at a 1:1 ratio to pre-diluted NanoLuc luciferase assay substrate. Samples were incubated in darkness for 5 minutes, before detection of luminescence by Molecular Devices ID5 plate reader, with an integration time of 10 s.

### 2.4 RNAP expression and purification

pET-29b-RNAP plasmids were transformed into BL21 or NEB Shuffle *E. coli* cells (New England Biolabs), and grown in 5 ml culture overnight at 37C. Starter cultures were used to inoculate 500 ml LB broth, which was incubated at 37 °C until an OD_600_ of 0.4 was reached. At this point incubation temperature was lowered to 25 °C. Protein expression was then induced by addition of 1M IPTG (Sigma-Aldrich), before harvesting of cells after 8 hours. Cell pellets were re-suspended for purification in buffer A (50 mM Tris-HCl, 0.5 M NaCl, pH 8.0), and lysed by sonication. After removal of cell debris by centrifugation (73000 x*g* for 15 mins), cell free extract was applied to a 5ml HisTrap HP column (Cytivia) at a flow rate of 5 ml/min. The column was washed with 2 column volumes Buffer A + 40 mM Imidazole (Sigma-Aldrich), before elution of protein in a gradient of imidazole from 0-300 mM over 10 column volumes. 5 ml of eluted protein was applied to a 1.6×60 cm Superdex200 gel filtration column at 1.5 ml/min, with 2 ml fractions collected after void volume. Fractions containing the RNAP of interest were concentrated to a final concentration of 1 mg/ml and exchanged into RNAP storage buffer (50 mM Tris-HCl, 100 mM NaCl, 50% v/v glycerol, 10 mM DTT, 0.1mM EDTA, 0.2% w/v NaN3 (All Sigma-Aldrich)), using a 50 kDA MWCO centrifugal filter (Sigma-Aldrich). RNAP preparations were stored at -20 °C. Purity of final preparations was assessed by Tris-Glycine SDS-PAGE 4-12% BT Novex gel with MES running buffer (Invitrogen).

### 2.5 In vitro transcription

Plasmid templates for IVT were linearised with XbaI, and purified by ethanol precipitation. Transcription reactions using the Hiscribe IVT kit (New England Biolabs), were assembled to a final volume of 20 μl. Reactions contained 2 μl 10X reaction buffer, 2 μl of each NTP, 1 ug of template DNA, and 2 μl of T7 RNAP, or 2 μl of novel RNAP. Transcription reactions were incubated for 2 hours, before addition of 1 μl DNase I, and further incubation for 20 minutes. Transcription reactions were purified using the Monarch RNA cleanup kit (New England Biolabs). mRNA concentration was quantified by nanodrop spectrophotometer, and product integrity assessed by agarose gel electrophoresis.

## 3. Results and Discussion

### 3.1 Bioinformatic analysis of the potential RNA polymerase biocatalyst solution space

The biocatalyst solution space for mRNA production is currently limited to a handful of characterised RNAPs. To define the theoretical solution space, we extracted the sequence of all putative single subunit RNAPs from Uniprot. At the time of conducting this analysis, 351 distinct single-stranded RNAPs had been predicted from publicly available genomics data. Accordingly, given that only six of these enzymes had been previously tested, approximately 98% of the potential solution space remained unexplored. We rationalized that determining the function of all 345 previously untested RNAPs would be highly-inefficient, and, moreover, unnecessary, given that many of these enzymes will share similar performance characteristics. Indeed, we assumed that variation in bioproduction phenotype (e.g., enzyme processivity, impurity generation profiles) would be underpinned by significant differences in amino acid sequences. Accordingly, we sought to define distinct spots within the potential solution space by identifying RNAP clusters that shared minimal amino acid sequence homology (Fig. 1A).

**Figure 1:**
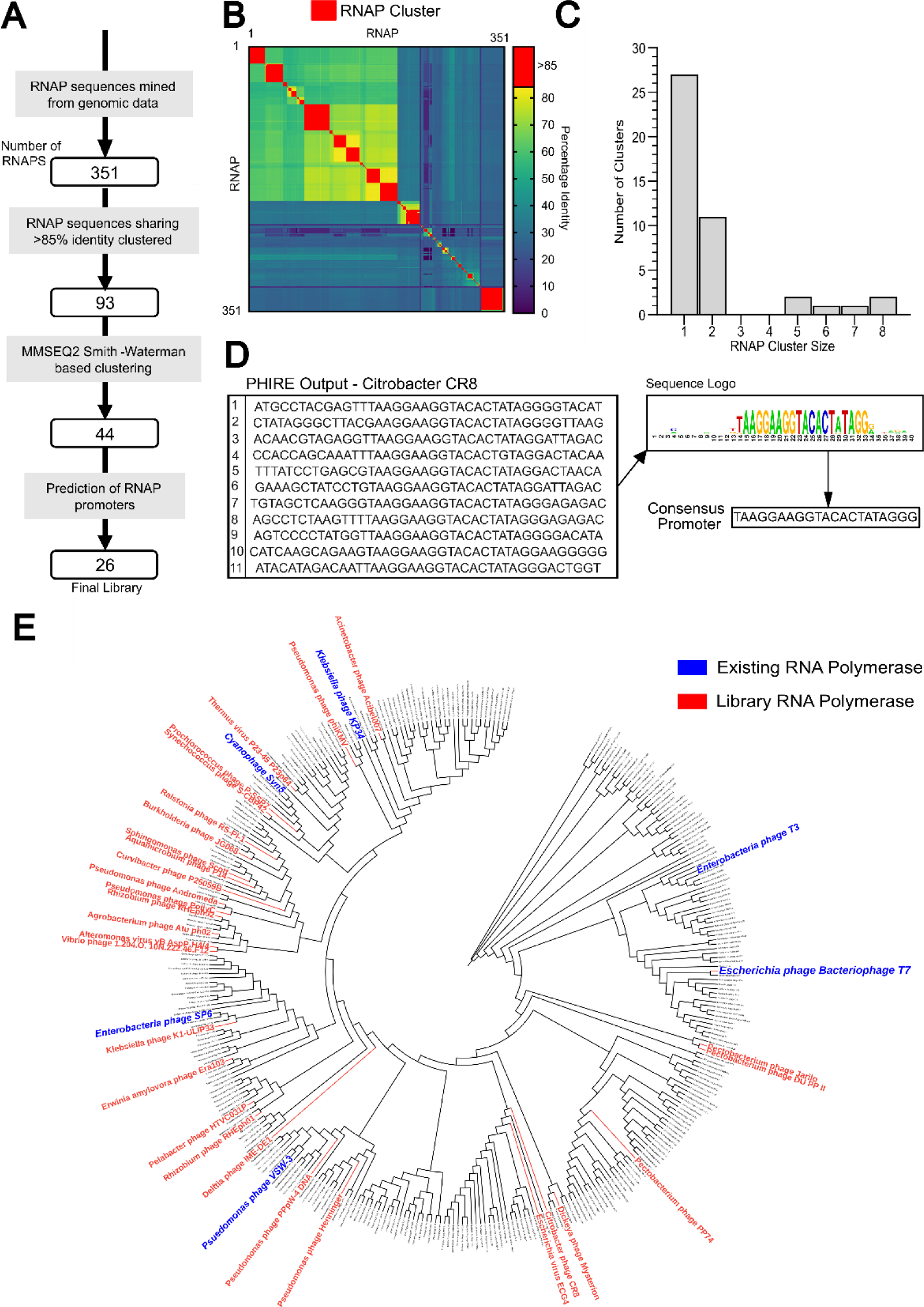
Bioinformatics-driven design of an RNAP ‘test’ library (**A**). Putative RNAPs were clustered using pairwise global sequence identity analysis (**B**), and subsequently grouped into 44 distinct families according to local sequence similarities (**C**). Representative polymerases from families for which accurate cognate promoter prediction was possible (**D**) were phylogenetically analysed to validate evolutionary diversity, represented in the circular cladogram showing all 351 analysed RNAPs (**E**).

The Clustal Omega sequence alignment tool (Madeira et al., 2019) was used to define RNAP clusters, whereby enzymes with >85% global sequence identity were grouped into a single distinct family. This analysis identified 93 enzyme clusters, where the smallest and largest groups contained 1 and 32 RNAPs respectively (Fig. 1B). To interrogate local sequence similarities, these families were then analysed using MMSEQ2, grouping RNAPs based on k-mer matching and the Smith-Waterman algorithm. Using sequence identity and coverage thresholds of 85% and 70% respectively (Steinegger and Söding, 2017), the number of discrete RNAP families was reduced from 93 to 44, where the majority of clusters (27) contained a single enzyme (Fig. 1C).

Utilisation of novel RNAPs for mRNA manufacture requires concomitant identification of appropriate cognate promoter elements to drive product transcription. This is non-trivial as single subunit RNAPs typically display highly stringent promoter binding activity, where mutation of a single nucleotide can abolish transcriptional output (Rong et al., 1998). Accordingly, identification of novel functional polymerases necessitates highly accurate promoter predictions. However, there are only limited publicly available tools to achieve this, where PHIRE (Lavigne et al., 2004) searches for conserved elements of defined length, and PhagePromoter utilises machine learning models to classify specified phage sequences as ‘promoter’ or ‘non-promoter’ (Sampaio et al., 2019). A representative polymerase was selected from each of the 44 families, and the associated phage genomes were investigated with both of these promoter prediction tools (Fig. 1D). This analysis failed to accurately identify cognate promoters in 18/44 cases. Testing further RNAPs from these 18 families similarly failed to result in identification of useable elements, indicating that the amino acid sequence diversity within these clusters is associated with ‘unusual’ cognate binding motifs that are significantly different to the promoter datasets that were used to train existing prediction tools. Accordingly, ∼40% of identified RNAP clusters could not be tested *in vitro* due to limitations in promoter prediction capabilities.

Cognate promoters were successfully predicted for the remaining 26 RNAPs, and optimal reaction temperatures were identified for each enzyme based on the growth temperature of corresponding phage hosts (Table 1). Ten RNAPs had predicted temperature optima ≤ 30°C, which may be beneficial for mRNA product quality profiles given that IVT reactions performed at reduced temperatures are associated with decreased levels of product-related impurities (Xia et al., 2022). Phylogenetic analysis of the 26 selected RNAPs confirmed that the final library comprised a panel of evolutionarily diverse enzymes, sharing no significant sequence similarity (≤ 75% global sequence identity) with any the six previously characterised polymerases (Fig. 1E). These polymerase-promoter pairs (Table 1) were taken forward for *in vitro* functional characterisation, facilitating testing of ∼60% of the identified RNAP clusters.

**Table 1:**
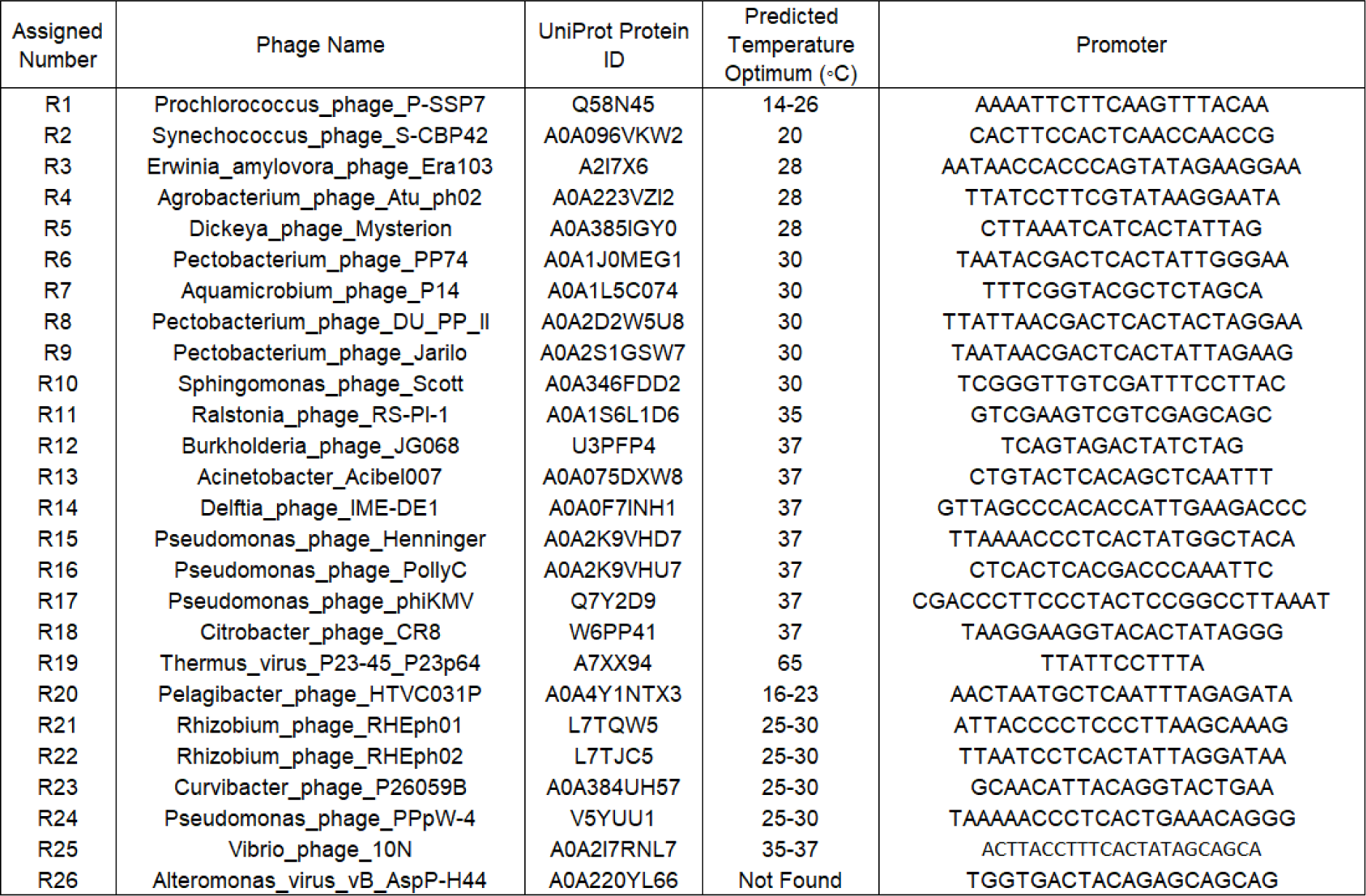
Bioinformatically-identified single-subunit RNA polymerases selected for *in vitro* functional characterization.

### 3.2 Identification of novel active RNAP biocatalysts via high-throughput *in vitro* functional characterisation

Previous studies focussed on identifying new RNAP biocatalysts for mRNA manufacture have relied on recombinant production of individual ‘test’ enzymes in *E. coli* cell-hosts, prior to characterisation in IVT reactions (Lu et al., 2019; Wang et al., 2022; Zhu et al., 2013). This time-consuming method is undesirable for characterisation of a large RNAP library, particularly given that manufacture of complex proteins at appropriate yield and quality can require significant process optimisation (Bhatwa et al., 2021; Gopal and Kumar, 2013). Moreover, variation in recombinant protein stability and purity may prevent accurate quantification of relative enzyme activities across the library. Accordingly, to functionally characterise our 26 novel RNAPs in parallel, we developed a high-throughput testing platform that does not require production and purification of each polymerase. This was achieved by adapting a cell-free coupled transcription-translation system that has previously been employed to rapidly assess activity of variant T7 polymerases (Egorova et al., 2021)(Cui et al., 2023). As shown in Figure 2A, this platform utilises a mastermix containing rabbit reticulocyte lysate and recombinant SP6 RNAP to facilitate *in vitro* production of a ‘test’ RNAP, which then in turn drives expression of a Nano-luciferase reporter-gene under the control of its cognate promoter.

**Figure 2:**
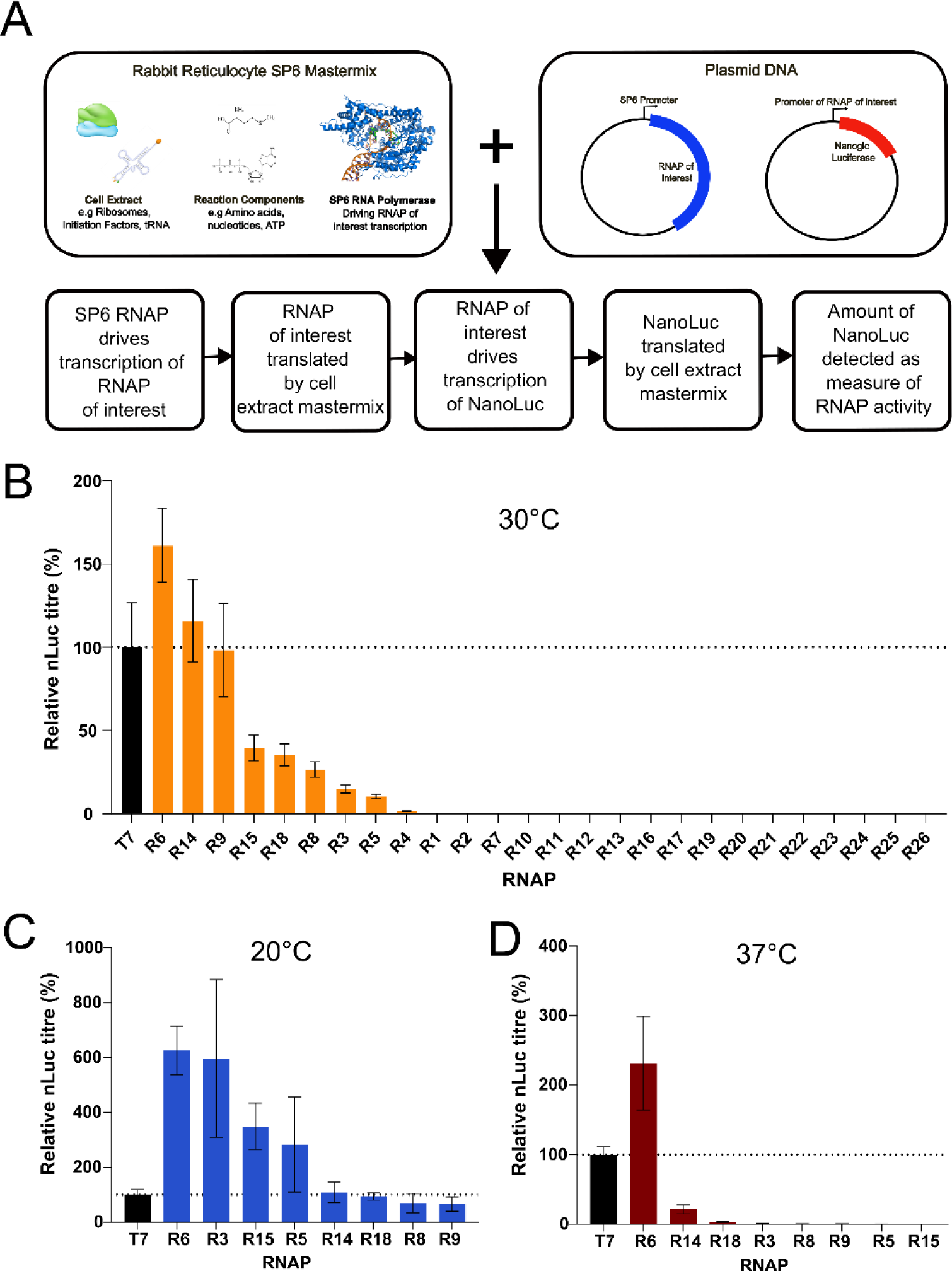
RNAPs were functionally characterised in a cell-free coupled transcription-translation system (**A**). Protein coding and cognate promoter sequence pairs were inserted into screening platform vectors and incubated with SP6-rabbit reticulocyte mastermix at 30°C (**B**), 20°C (**C**) and 37°C (**D**). Luciferase expression was quantified 1 hr post-incubation; data are expressed as a percentage of the production achieved using the control T7 RNAP. Values represent the mean + *SD* of three independent experiments (*n* = 3, each performed in triplicate).

Protein coding sequences and predicted cognate promoter elements for each of the 26 test enzymes were chemically synthesized and inserted into the appropriate RNAP screening platform vectors (Fig. 2A). Resulting plasmid-pairs were individually mixed with the SP6 RNAP-rabbit reticulocyte lysate mastermix, and Luciferase production was measured after incubating the reaction for 1 hr at 30°C (recommended assay reaction temperature). As shown in Figure 2B, 8/26 enzymes were functionally active, driving luciferase expression levels that ranged from 10% - 161% of that achieved using the control T7 RNAP. Accordingly, approximately 70% of tested enzymes were non-functional, highlighting the difficulty associated with identifying novel RNAP biocatalysts.

There was no significant correlation between predicted enzyme temperature optima and observed activity at 30°C. However, to further assess the impact of reaction parameters on polymerase performance, we tested enzyme activities at increased (37°C) and decreased (20°C) temperatures. While the same eight RNAPs were functional at 20°C, only three of these enzymes displayed activity at 37°C. Moreover, apart from R6 which drove highest Luciferase expression levels under all conditions tested, the relative performance of polymerases varied with temperature. Although our objective was to identify effective polymerase-promoter pairs, rather than to precisely elucidate their relative performance characteristics, these data indicate that enzymes active over a narrow range of temperatures may be incorrectly categorized as non-functional. However, we concluded that this was unlikely when testing across three separate temperature set-points, and that enzyme inactivity in our screening platform was more likely due to either i) inaccurate annotation/sequencing of putative RNAP coding sequences or ii) incorrect promoter prediction.

Although enzyme activity in the cell-free screening system may not be directly predictive of performance in IVT-based mRNA manufacturing processes, it is notable that polymerase R6 drove higher levels of luciferase expression than T7 in all conditions tested (increase ranging between 160% – 620%), including a 220% increase at T7s optimum reaction temperature (37°C). Five further RNAPs (R3, R5, R9, R14, R15) facilitated luciferase titres greater than or equal to that achieved with T7 in at least one reaction condition. Accordingly, these enzymes may exhibit higher processivity/catalytic activity than T7 and could therefore have potential use in enhancing mRNA production yields. Moreover, their use may permit simplified downstream processing operations via reduced formation of product-related impurities, particularly as many of these RNAPs exhibit relatively high activities at low temperatures (Wang et al., 2022). While further characterisation is required to fully assess their bioindustrial utility, the identification of eight novel functional enzymes more than doubles the number of available RNAPs, expanding the biocatalyst solution space for mRNA manufacture by ∼130%.

### 3.3 Cognate Promoter prediction is the critical limiting factor restricting further expansion of the RNAP biocatalyst solution space

The finding that ∼70% of characterized enzymes were non-functional in *in vitro* tests (Fig. 2) indicates that the vast majority of putative RNAPs cannot be simply extracted from online databases and directly employed in mRNA manufacturing applications. Given that RNAPs are known to display highly stringent promoter recognition requirements (Rong et al., 1998), we hypothesised that enzyme inactivity may have resulted from inaccurate predictions of cognate promoter sequences. To exemplify this, we characterized the ability of R6, the best performing polymerase in *in vitro* screens, to initiate transcription from the promoters of other functional enzymes. As shown in Figure 3A, R6 could not drive quantifiable gene expression from any of these variant elements, where even a single nucleotide change was sufficient to completely abolish transcriptional output. These data highlight that the ability to exploit any given potential RNAP biocatalyst is heavily dependent on highly accurate definition of its cognate promoter sequence.

**Figure 3:**
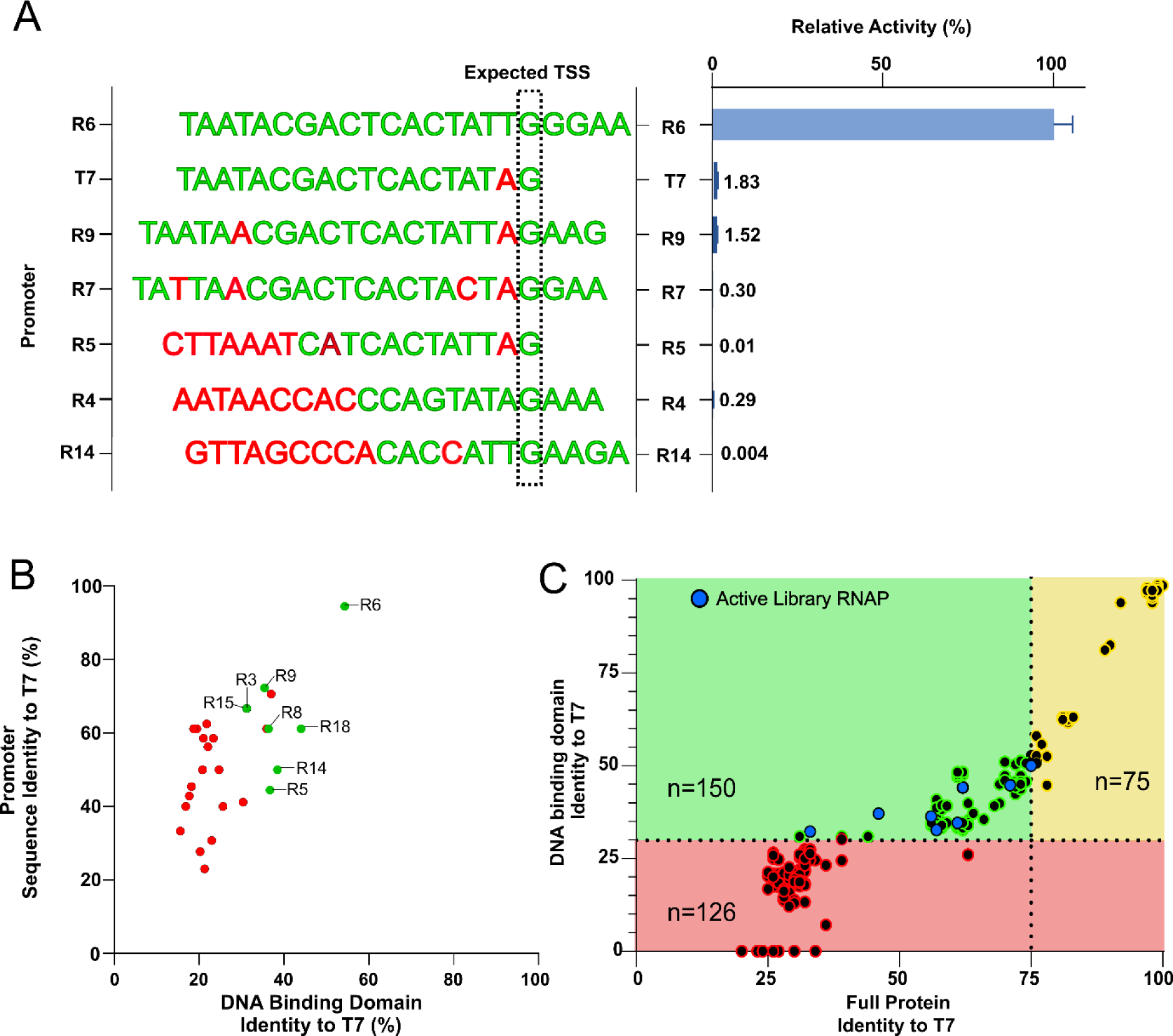
**A**) The ability of RNAP6 (Pectobacterium phage PP74) to drive Luciferase expression from varying non-cognate promoter elements was evaluated in cell-free coupled transcription-translation assays (see Fig. 2). Data are expressed as a percentage of the production achieved using the cognate RNAP-6 promoter. Values represent the mean + SD of three independent experiments (*n* = 3, each performed in triplicate). **B**) Test library RNAP-promoter pairs were analysed to determine relative DNA binding domain and promoter sequence identity with T7. Pairs that were found to be active or inactive in functional characterisation tests are shown as green and red dots respectively. **C**) The entire theoretical RNAP biocatalyst solution space was analysed to identify promising future targets for *in vitro* characterisation (green section). Enzymes predicted to have incorrect promoter definitions or similar bioproduction phenotypes to T7 are shown in the red and yellow sections respectively. Functional RNAPs identified in this study are shown as blue dots.

We reasoned that RNAP promoter prediction tools may be incapable of precisely defining new elements that are significantly divergent from currently known sequences, as evidenced by our inability to derive cognate promoters for ∼40% of bioinformatically-determined RNAP clusters (see Section 3.1). Indeed, given the paucity of characterised RNAP promoters, novel ‘test’ enzymes may recognise sequence motifs and architectures that are i) substantially different to those used to train/design current algorithms, and accordingly ii) beyond the predictive capabilities of available tools. Rationalising that divergence in promoter structure/sequence would be underpinned by differences at the amino acid level, we investigated whether enzyme inactivity was associated with DNA binding domain sequences that varied significantly to those of well-studied biocatalysts. As shown in Figure 3B, 8/10 polymerases that share relatively high DNA binding domain sequence identity with T7 (>30%) were found to be active, while all 16 enzymes that share relatively low similarity (<30%) were non-functional. In contrast, cognate promoter sequence similarity with T7 promoter was not a good predictor of RNAP functionality, where 6/8 active and 12/16 inactive elements shared between 40% and 65% sequence identity with T7 (Fig. 2B). These data suggest that we lack the ability to accurately predict divergent promoter elements for new RNAPs when shared DNA binding domain sequence identity with well-studied enzymes falls below a critical threshold.

Our findings indicate that potential RNAPs can be efficiently screened *in silico*, where enzymes that share below ∼30% DNA-binding domain sequence identity with T7 are unlikely to be functional *in vitro* owing to incorrect promoter definition. However, as shown in Figure 3C, this cut-off removes approximately 36% of the theoretical biocatalyst solution space for mRNA production. Of the remaining 225 polymerases, 75 share relatively high overall protein sequence identity (>75%) with T7. Such enzymes are considered unlikely to exhibit substantial differences to T7 in key performance criteria such as enzyme processivity and product-related impurity generation. Accordingly, only 150 RNAPs are predicted to be both active *in vitro* and potentially display novel, desirable bioproduction functionalities (including the eight we have identified in this study). This analysis therefore highlights 142 promising additional biocatalyst targets for future investigation, including 88 that do not share high sequence identity (>75%) with either the 6 previously characterized RNAPs or the 8 enzymes identified in this study (listed in Supplementary table 1). However, it also suggests that >120 potentially useful enzymes, are currently difficult to exploit, highlighting promoter prediction capability as the key limiting factor preventing comprehensive exploitation of the theoretical biocatalyst solution space for mRNA production. Although the cognate promoters of individual polymerases can be elucidated via non-bioinformatic laboratory techniques (Lu et al., 2019), these time-intensive methods are intractable when testing multiple enzymes in parallel. Accordingly, full exploration of the putative RNAP biocatalyst solution space to optimise mRNA production processes will likely require significant advancements in phage promoter prediction tools.

### 3.4 Novel identified RNAPs enhance the biocatalyst solution space for IVT-based mRNA production

To validate that novel RNAPs identified via our HT cell-free screening platform have utility in mRNA manufacturing processes, we recombinantly produced polymerases R5 and R6 in *E. coli*. These polymerases were chosen to represent highly- and moderately-active enzymes, where R6 (Pectobacterium phage PP74) was previously shown to be the best performing RNAP in all temperatures tested, and R5 (Dickeya phage Mysterion) drove relatively low-to-medium levels of transcription across varying reaction conditions (Fig 2). Polymerases were overexpressed in 0.5 L scale production processes and purified using His-tag affinity and size exclusion chromatographic operations. Purified recombinant RNAPs were then utilised in IVT reactions to manufacture *Cypridina* Luciferase (CLuc) mRNA. As shown in Figure 4, both enzymes drove significant levels of Cluc expression, validating their function as biocatalysts for synthetic mRNA production. To evaluate enzyme robustness, we tested the performance of each RNAP at a range of pH (predicted optimum ± 1) and temperature (predicted optimum ± 5°C) set-points. Both RNAPs were functional across all conditions tested, where R5 performance was relatively constant, and R6 activity increased with temperature. The latter highlights that expected phage host growth temperatures are not directly predictive of optimal *in vitro* reaction conditions for recombinant RNAPs.

**Figure 4:**
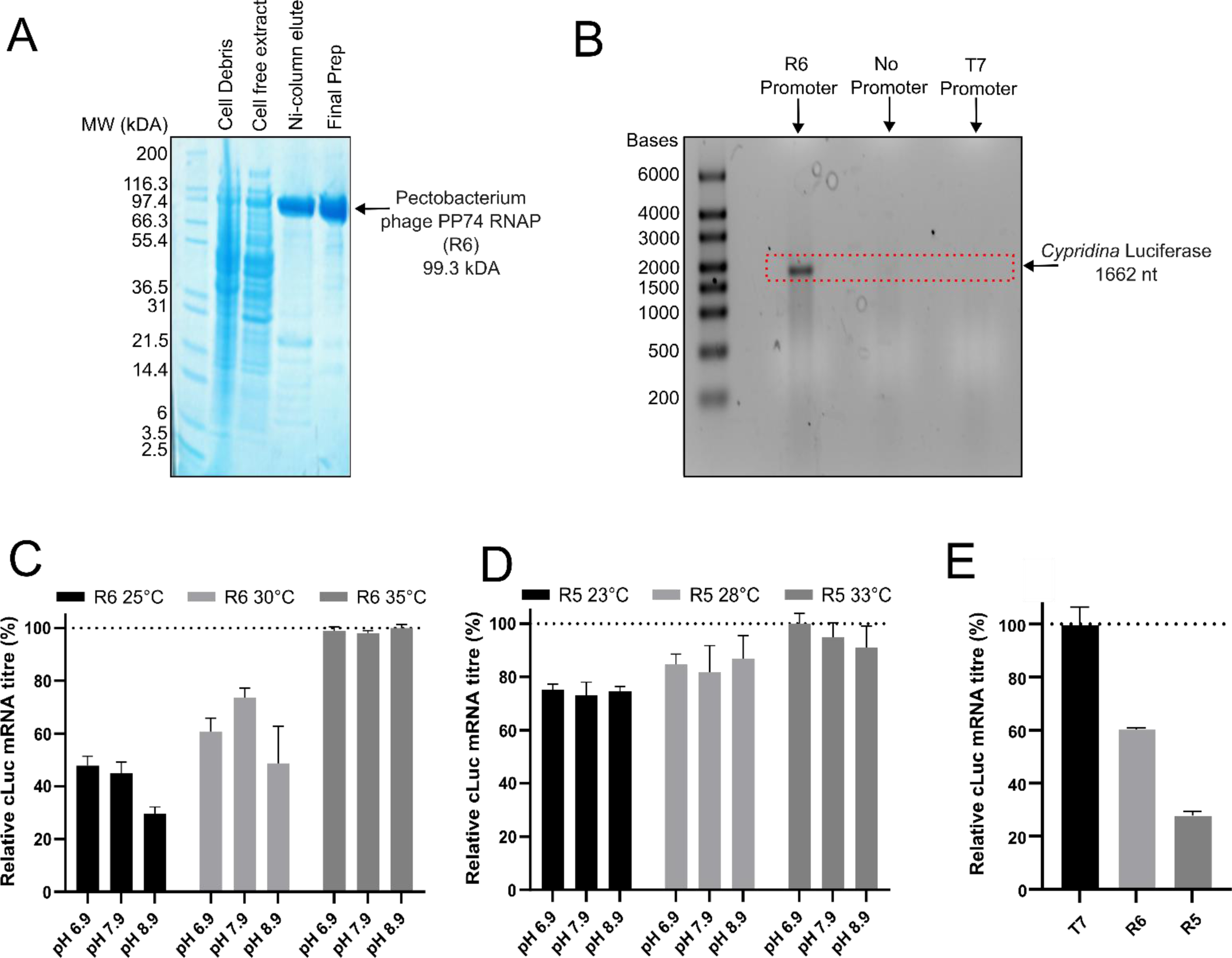
RNAP 5 and 6 were recombinantly produced in *E. coli* and purified using His-tag affinity and size exclusion chromatography (**A**). Purified enzymes were used to manufacture Luciferase mRNA in *in vitro* transcription (IVT) reactions, and production of full-length product was verified by gel electrophoresis analysis (**B**). IVT production processes utilising RNAP6 (**C**) and RNAP5 (**D**) were performed in varying reaction conditions and resulting mRNA titers were quantified using nanodrop spectrophotometry. Using identified optimal reaction parameters for each enzyme, the relative performance of recombinant polymerases was evaluated compared to an NEB T7 control (**E**). Data in C and D are expressed as a percentage of the production achieved using optimal reaction parameters for each enzyme. Data in E are expressed as a percentage of the production achieved using T7. In C, D and E values represent the mean + SD of three independent experiments (*n* = 3, each performed in triplicate).

As shown in Figure 4, utilisation of R6 at ‘optimal’ reaction parameters (pH 7.9, 35°C) facilitated mRNA product titers >60% of that achieved when using NEB recombinant T7 at recommended conditions (pH 7.9, 37°C). Although R6 drove higher levels of gene transcription than T7 in our cell-free system, it is not surprising that T7s relative activity was enhanced in IVT processes given that NEB T7 is a highly-pure engineered enzyme with fully-optimised reaction conditions. Indeed, we anticipate that rational protein engineering/evolution, coupled with improved purification techniques and reaction parameters (E.g., optimised MgCl_2_ concentration), will significantly increase R6s biocatalytic activity in IVT-based mRNA production. Irrespective of this, by initially facilitating product yields equivalent to ∼61% of that achieved by optimised T7, R6 is considered a highly active biocatalyst.

Polymerase R5 enabled product titers ∼47% of that achieved using R6, suggesting that the comparative performance of novel RNAPs in cell-free testing platforms is broadly predictive of their relative ability to maximise mRNA yields in IVT manufacturing processes. Accordingly, we concluded that the additional 6 novel enzymes identified in this study (Erwinia amylovora phage Era103, Pectobacterium phage DUPP II, Pectobacterium phage Jarilo, Delftia phage IME-DE1, Pseudomonas phage Henninger, Citrobacter phage CR8) are also likely to facilitate moderate-to-high mRNA production yields. While we cannot currently comment on the relative ability of these new polymerase to enhance product quality, previous work suggests they will generate variable levels of product-related impurities, such as dsRNA and truncated species (Lu et al., 2019; Wang et al., 2022; Xia et al., 2022; Zhu et al., 2013, 2015). Indeed, this new library is particularly likely to exhibit differential bioproduction phenotypes, given that they were specifically selected based on sharing minimal amino acid sequence similarities. We therefore conclude that addition of these eight novel functional enzymes to the RNAP biocatalyst solution space will significantly enhance IVT-based mRNA manufacturing optimisation strategies.

## 4. Concluding Remarks

The eight novel sequence-diverse functional RNAPs identified in this study substantially increases the number of biocatalysts available for mRNA production. Although further work is required to comprehensively define their relative performance characteristics, particularly their associated impurity generation profiles, this expansion of the biocatalyst solution space significantly enhances design options for molecule-, process-, and application-specific optimisation of mRNA product yield and quality. This improved flexibility will become increasingly useful as mRNA product lines continue to diversify towards large, complex molecules that pose new manufacturing challenges (Bai et al., 2023; Blakney et al., 2021; Chen et al., 2022; Pourseif, 2022; Qu et al., 2022). Our combined *in silico* and *in vitro* analysis of the theoretical biocatalyst solution space i) showed that full exploitation of potential RNAPs is restricted by cognate promoter prediction capabilities, but, also ii) identified a panel of enzymes that are particularly promising for future investigation. With respect to the latter, the screening platform we have developed can be utilised to rapidly test additional RNAP libraries, including engineered variants of existing polymerases. In conclusion, by more than doubling the number of available polymerases, and providing associated methods to select and screen further new enzymes, this study has facilitated a significant expansion of the RNAP biocatalyst solution space, enhancing strategies to optimise and troubleshoot IVT-based mRNA production processes.

## Author Contributions

**Edward Curry** – Conceptualisation, methodology, analysis, investigation, manuscript preparation. **Svetlana Sedelnikova** – Investigation, analysis. **John Rafferty** – Investigation, analysis. **Martyn Hulley** – Conceptualisation, analysis. **Adam Brown** – Conceptualisation, methodology, analysis, manuscript preparation, funding acquisition.

## Data sharing statement

The data that support the findings of this study are available from the corresponding author upon reasonable request.

## Conflict of interest statement

The authors declare no conflict of interest

## Funding Statement

This study was supported by AstraZeneca and the Biotechnology and Biological Sciences Research Council (Grant No. BB/T508664/1)

## Abbreviations

RNAP: RNA Polymerase
IVT: *In vitro* transcription
CLuc: Cypridina Luciferase
dsRNA: Double stranded RNA

## Notes

### Competing Interest Statement

The authors have declared no competing interest.

