## Supplementary Table 1 for "Expanding the RNA polymerase biocatalyst solution space for mRNA manufacture"

**Supplementary Table 1**: RNAPs identified as promising targets for future *in vitro* characterisation studies based on having i) relatively high (>30%) DNA binding domain identity with T7, and ii) relatively low (< 75%) overall sequence identity with currently available polymerases.

| **RNAP Name** | **Identity to T7 (%)** | **DNA binding domain identity (%)** | **Uniprot ID** |
| --- | --- | --- | --- |
| Dickeya phage Ninurta | 70 | 47.33 | A0A2S1GTB4 |
| Dickeya phage vB_DsoP_JA10 | 70 | 46 | A0A384ZVU4 |
| Enterobacteria phage K11 | 72 | 42.38 | P18147 |
| Erwinia phage pEp_SNUABM_09 | 73 | 47.33 | A0A5J6DAB1 |
| Erwinia phage vB_EamP-L1 | 69 | 45.03 | G0YQ47 |
| Escherichia phage K30 | 74 | 45.7 | F8R4Q2 |
| Escherichia phage Pisces | 62 | 32.68 | A0A5B9NC24 |
| Klebsiella phage 2044-307w | 73 | 45.03 | A0A249Y210 |
| Klebsiella phage IME304 | 74 | 45.7 | A0A4Y5TVL6 |
| Klebsiella phage K11 | 72 | 45.03 | B3VCY2 |
| Klebsiella phage K5 | 72 | 45.03 | A0A0F7LBY1 |
| Klebsiella phage K5-4 | 73 | 45.7 | A0A219YHD6 |
| Klebsiella phage KN1-1 | 73 | 45.03 | A0A3S5IBH2 |
| Klebsiella phage KN3-1 | 73 | 45.03 | A0A3Q9WWY9 |
| Klebsiella phage KN4-1 | 73 | 45.03 | A0A3Q9WSE6 |
| Klebsiella phage KOX3 | 73 | 45.7 | A0A5B9NGK0 |
| Klebsiella phage KOX5 | 73 | 45.7 | A0A5B9NDA1 |
| Klebsiella phage KP32 | 72 | 45.7 | D1L2T7 |
| Klebsiella phage kpssk3 | 73 | 45.03 | A0A3G8F354 |
| Klebsiella phage Kund-ULIP47 | 73 | 45.7 | A0A4P6DBN7 |
| Klebsiella phage Kund-ULIP54 | 73 | 45.7 | A0A4P6PMA0 |
| Klebsiella phage Pharr | 73 | 45.03 | A0A4D6DY13 |
| Klebsiella phage SH-Kp 152410 | 73 | 44.37 | A0A2K9VGQ6 |
| Klebsiella phage vB_Kp1 | 72 | 45.03 | A0A0P0IV82 |
| Klebsiella phage vB_KpnP_BIS33 | 73 | 43.71 | A0A1V0E6J1 |
| Klebsiella phage vB_KpnP_IME205 | 74 | 45.7 | A0A0U3DFB5 |
| Klebsiella phage vB_KpnP_IME321 | 73 | 45.7 | A0A344UBX8 |
| Klebsiella phage vB_KpnP_IME335 | 73 | 45.03 | A0A5J6CUK8 |
| Klebsiella phage vB_KpnP_KpV289 | 72 | 44.37 | A0A0K2YWK9 |
| Klebsiella phage vB_KpnP_KpV763 | 73 | 45.03 | A0A1D8F0C6 |
| Klebsiella phage vB_KpnP_KpV766 | 73 | 45.03 | A0A1I9SFA1 |
| Klebsiella phage vB_KpnP_KpV767 | 73 | 45.7 | A0A1I9SF50 |
| Klebsiella phage vB_KpnP_NahiliMali | 70 | 47.33 | A0A5B9NQY1 |
| Klebsiella phage vB_KpnP_PRA33 | 73 | 45.7 | A0A1V0E683 |
| Klebsiella phage vB_KpnP_Sibilus | 70 | 46 | A0A5B9NKC7 |
| Morganella phage MmP1 | 70 | 51.02 | D1FNQ5 |
| Morganella phage vB_MmoP_MP2 | 72 | 50.34 | A0A192YBW9 |
| Pectobacterium phage DU_PP_II | 64 | 34.57 | A0A2D2W5U8 |
| Pseudomonas phage 22PfluR64PP | 57 | 35.71 | A0A3G6V715 |
| Pseudomonas phage 67PfluR64PP | 57 | 35.71 | A0A2S1PGT5 |
| Pseudomonas phage 71PfluR64PP | 57 | 35.71 | A0A2S1PDT9 |
| Pseudomonas phage Pf-10 | 58 | 38.31 | A0A0A0YSI2 |
| Pseudomonas phage PFP1 | 57 | 36.36 | A0A2Z4QIP2 |
| Pseudomonas phage phi15 | 57 | 40.91 | F0V6X0 |
| Pseudomonas phage Phi-S1 | 58 | 38.31 | M4H3N8 |
| Pseudomonas phage PPPL-1 | 57 | 38.71 | A0A0S2MVL3 |
| Pseudomonas phage PspYZU08 | 56 | 34.64 | A0A2U7NJN9 |
| Pseudomonas phage shl2 | 58 | 39.35 | A0A160SY77 |
| Ralstonia phage DU_RP_I | 39 | 30.94 | A0A2D2W578 |
| Ralstonia phage RSB1 | 31 | 30.94 | A0A5P8D3T3 |
| Ralstonia phage RSB2 DNA | 44 | 30.94 | A0A5P8D447 |
| Ralstonia phage RsoP1EGY | 39 | 30.22 | A0A2R2ZGE5 |
| Vibrio phage ICP3 | 62 | 48.32 | F1D002 |
| Vibrio phage ICP3_2007_A | 62 | 48.32 | F1D0J5 |
| Vibrio phage ICP3_2008_A | 61 | 48.32 | F1D0E7 |
| Vibrio phage ICP3_2009_A | 62 | 48.32 | F1D0A0 |
| Vibrio phage ICP3_2009_B | 62 | 47.65 | F1D053 |
| Vibrio phage JSF11 | 62 | 48.32 | A0A2D0Z112 |
| Vibrio phage JSF18 | 62 | 48.32 | A0A2D0YMX9 |
| Vibrio phage JSF20 | 62 | 48.32 | A0A2D0YL99 |
| Vibrio phage JSF24 | 62 | 48.32 | A0A2D0Z841 |
| Vibrio phage JSF25 | 62 | 48.32 | A0A2D0XR33 |
| Vibrio phage JSF30 | 62 | 48.32 | A0A2D0YV87 |
| Vibrio phage JSF31 | 61 | 46.98 | A0A2D0Z1P3 |
| Vibrio phage JSF32 | 61 | 46.98 | A0A2D0Z2L0 |
| Vibrio phage JSF34 | 62 | 48.32 | A0A2D0YLK2 |
| Vibrio phage JSF35 | 62 | 47.65 | A0A2D0YKN7 |
| Vibrio phage JSF36 | 61 | 47.65 | A0A2D0Z259 |
| Vibrio phage N4 | 61 | 48.32 | D0Q187 |
| Vibrio phage Rostov-1 | 62 | 48.32 | A0A2P0ZKC2 |
| Vibrio phage VP3 | 62 | 48.32 | H9YAF4 |
| Vibriophage VP4 | 61 | 48.32 | Q4TVY1 |
| Yersinia phage fPS-10 | 73 | 51.33 | A0A2H1X8U9 |
| Yersinia phage fPS16 | 73 | 51.33 | A0A2H1X8Z7 |
| Yersinia phage fPS-19 | 73 | 51.33 | A0A2D0PDM8 |
| Yersinia phage fPS-21 | 73 | 51.33 | A0A2D0PE32 |
| Yersinia phage fPS-26 | 73 | 51.33 | A0A2D0PD60 |
| Yersinia phage fPS-50 | 73 | 51.33 | A0A2D0PDY6 |
| Yersinia phage fPS-52 | 73 | 51.33 | A0A2D0PDI2 |
| Yersinia phage fPS-53 | 73 | 51.33 | A0A2H1UJD0 |
| Yersinia phage fPS-54-ocr | 73 | 51.33 | A0A2H1UJE6 |
| Yersinia phage fPS-59 | 73 | 51.33 | A0A2D0PE84 |
| Yersinia phage fPS-64 | 73 | 51.33 | A0A2D0PEF0 |
| Yersinia phage fPS-7 | 73 | 51.33 | A0A2D0PDD4 |
| Yersinia phage fPS-85 | 73 | 51.33 | A0A2H1UKL9 |
| Yersinia phage fPS-86 | 73 | 51.33 | A0A2D0PEP1 |
| Yersinia phage fPS-89 | 73 | 51.33 | A0A2D0PDP8 |
| Yersinia phage fPS-9 | 73 | 51.33 | A0A2C9D120 |
